# CRISPR-Cas9 precision editing of kinetochore protein phosphosite codons in *Leishmania mexicana*

**DOI:** 10.64898/2026.01.15.699675

**Authors:** Charlotte McNiven, Juliana B. T. Carnielli, Vincent Geoghegan, Joana R.C. Faria, Jeremy C. Mottram

**Affiliations:** York Biomedical Research Institute and Department of Biology, University of York, UK

## Abstract

*Leishmania mexicana*, like other trypanosomatids, possesses a unique kinetochore—the protein complex crucial for chromosome segregation during mitosis. To investigate the functional significance of specific phosphorylation sites on essential kinetochore proteins, we adapted a selection-free precision editing strategy using CRISPR-Cas9 in *Leishmania mexicana* promastigotes. Our method targeted genomic DNA with 160-bp double-stranded DNA repair templates and guide RNAs to introduce targeted modifications. We focused on six phosphosites within the kinetochore proteins KKT2, KKT4, and KKT7, generating phosphodeficient, phosphomimetic, and synonymous mutants for each site. Across 18 independent transfections, we achieved a successful editing rate of 27.5% as determined by PCR screening, with 30.4% of clones confirmed as edited by Sanger sequencing. A significant portion of these edited clones (22.1%) were homozygous. Despite these precise genomic modifications, none of the phosphosite mutant clones exhibited any apparent growth defects or cell cycle dysregulation, suggesting these phosphorylation sites individually may not be critical for these processes under standard culture conditions. To facilitate higher-throughput precision editing, we developed a Python script that automates the design of the 160-bp repair templates. This script uses a FASTA file, a codon usage table, and a simple configuration file to design templates with a single nonsynonymous mutation and additional synonymous mutations for screening purposes. It also generates a corresponding synonymous-only repair template and primers for both screening and repair template generation, offering a “ready-to-go” approach. While designed for *Leishmania*, this powerful tool is adaptable for use with other kinetoplastids.

## Introduction

*Leishmania* species, the causative agents of leishmaniasis, are evolutionarily distinct eukaryotic parasites with highly unusual and divergent genomes. Understanding the function of the unique proteins encoded by these genomes is critical for developing new therapeutics. For decades, gene editing has been a key investigative approach (Cruz and Beverley, 1990), a field that was revolutionized in *Leishmania* by the adaptation of the CRISPR-Cas9 system (Sollelis et al., 2015; Beneke et al., 2017). The principle of Cas9-mediated editing is elegant: a guide RNA directs the Cas9 endonuclease to a specific genomic locus, creating a double-stranded DNA break (DSB). The cell’s natural DNA repair machinery, specifically the Homology Directed Repair (HDR) pathway, is then “hijacked” to incorporate a synthetic DNA template. Critically, initial assumptions that HDR in *Leishmania* required kilobases of homologous DNA have been dispelled, as it has since been shown that as little as 30 base pairs (bp) of homology are sufficient when a Cas9-induced DSB is present (Beneke et al., 2017).

Despite this technical advance, most established Cas9-based methods in *Leishmania* still rely on a positive selection marker to isolate successfully modified cells. While effective, this reliance imposes significant limitations. The large constructs required restrict modifications primarily to gene termini (e.g., epitope tagging) or full gene deletions. Introducing subtle, single-codon changes—such as those needed to study post-translational modifications—becomes a laborious process. It typically requires *in vitro* synthesis of a new gene copy and subsequent replacement of the native one, a time-consuming method susceptible to disruption of endogenous protein expression and regulation (Boucher et al., 2002; da Silva et al., 2012; Terrão et al., 2017; Tupperwar et al., 2019; Borges et al., 2024; Ribeiro et al., 2024). This drawback highlights the urgent need for a high-efficiency method of precision editing that uses much smaller, marker-free repair templates. Though this approach has been successful in a range of organisms, including other kinetoplastids, an efficient and consistent selection-independent protocol for *Leishmania* remains a technical bottleneck (Zhang and Matlashewski, 2015; Rico et al., 2018; Wall et al., 2018; Altmann et al., 2022; Carnielli et al., 2025a, 2025b; Lansink et al., 2025; Novotná et al., 2025).

A crucial, yet poorly understood, target for this precise genetic analysis is the kinetochore. This essential protein complex links chromosomal DNA to the mitotic spindle, ensuring accurate segregation of sister chromatids during mitosis (Musacchio and Desai, 2017). In stark contrast to canonical eukaryotes, the kinetochore of kinetoplastids is highly divergent, with most of its components lacking homology to well-characterized proteins (Akiyoshi and Gull, 2014). To date, 25 unique Kinetoplastid Kinetochore proteins (KKTs) and 12 interacting proteins (KKIPs) have been identified in *Trypanosoma brucei*, which are largely conserved in *Leishmania* (Akiyoshi and Gull, 2014; Nerusheva and Akiyoshi, 2016; Nerusheva et al., 2019; Geoghegan et al., 2022). Our structural understanding of this machinery is rudimentary, but components like the inner kinetochore kinases KKT2 and KKT3 are known to be essential for viability and assembly of the kinetochore complex in *Leishmania* (Baker et al., 2021; Geoghegan et al., 2022).

Recent phosphoproteomic studies have revealed that protein phosphorylation is a key dynamic regulator of kinetochore function in *Leishmania mexicana* (Geoghegan et al., 2022). Specifically, phosphorylation of proteins such as KKT2, KKT4, and KKT7 changes significantly throughout the cell cycle, suggesting these post-translational modifications play a crucial role in controlling mitotic progression. However, both the precise function of individual phosphorylation sites and the kinases responsible for them are largely unknown. Isolating the effect of a single amino acid change in this complex system requires an efficient, high-throughput precision editing strategy.

In this study, we addressed the technical and biological gaps by developing and optimizing an efficient, selection-independent precision editing methodology for *Leishmania mexicana*. We generated phosphodeficient, phosphomimetic, and synonymous mutants at six distinct phosphosites in KKT2, KKT4, and KKT7 to determine the functional role of these individual sites. Our optimized method, which utilizes double-stranded DNA repair templates with 50 bp homology arms, achieved a high efficiency, with 22.1% of screened clones being homozygous precision mutants. This result represents a significant improvement over single-stranded oligonucleotide-based approaches, which overall only generated 2.8% edited clones. The single phosphosite mutants we generated lacked a distinct growth or cell cycle phenotype, suggesting that phosphorylation-mediated regulation of these essential proteins may be redundant or context-dependent. Finally, to support future precision editing efforts in *L. mexicana* and other kinetoplastids, we developed a Python script that automates the design of all necessary oligonucleotides for both repair template production and PCR-based screening. This tool provides a streamlined, ready-to-use platform to enable further high-resolution genetic analysis in these divergent organisms.

## Results

### Single-stranded oligonucleotides generate selection-free precision edited kinetochore phosphosite mutants at low efficiency in Leishmania mexicana

Previous reports of CRISPR-Cas9-mediated precision editing in kinetoplastids primarily relied on single-stranded oligodeoxynucleotide (ssODNs) as repair templates. Crucially, most of these successful studies utilized mutations that conferred drug resistance, allowing for strong selection and enrichment of edited clones (Zhang and Matlashewski, 2015; Medeiros et al., 2017; Rico et al., 2018; Carnielli et al., 2025a).

To develop a selection-free protocol, we first tested the ssODN strategy in *Leishmania mexicana* T7Cas9 cells. We designed 120 nt ssODN templates with 30 nucleotide homology arms to target specific phosphosites on the kinetochore proteins KKT2, KKT4, and KKT7. We selected phosphosites on these proteins that were conserved across a range of *Leishmania* species and were shown by spatial phosphoproteomics to be proximal to KKT3 (Geoghegan et al., 2022). Many of the sites we chose were dynamically altered during the cell cycle, were affected by CLK1 kinetochore protein kinase inhibition or were near domains predicted to be functionally important (see Supplementary Table 1 for more details) (Geoghegan et al., 2022). To prevent the active Cas9 enzyme from re-cutting the edited locus, we strategically recoded both the protospacer and PAM (protospacer adjacent motif) sequences along with the target codon, a modification not commonly implemented in published ssODN protocols.

Despite screening 107 clones across multiple sites, this approach showed low efficiency for generating phosphosite mutants on kinetochore proteins. We recovered only three mutant clones (2.8% of the total screened), with only two of these being homozygous integrations (Table 1). For example, transfections targeting KKT7 S304A yielded zero mutants out of the 20 clones screened, and the most successful target, KKT2 S493A, only produced 3 edited clones out of 41. No mutant clones were recovered from the KKT7 S304S synonymous mutant control transfection. The failure to recover clones with synonymous mutations points towards suboptimal editing efficiency as the limiting factor, although synonymous recoding has been reported to affect protein expression in trypanosomatids (Jeacock et al., 2018; Nascimento et al., 2018), which may also account for the low recovery rate observed. Collectively, these findings indicate that ssODN-mediated editing represents a labour-intensive approach for selection-free precision editing of phosphosites in *Leishmania* kinetochore proteins. This poor recovery rate demonstrated that, when the mutation being induced is not selectable (e.g. does lead to drug-resistance), use of ssODNs is not an effective strategy for efficient precision editing in *L. mexicana*.

**Table 1.** Number of mutant clones recovered from transfections using single-stranded DNA repair templates.

| *Target mutation* | *Clones assessed* | *Homozygous mutants* | *Heterozygous mutants* | *Complex mutants<sup>‡</sup>* | *Total Mutants* |
| --- | --- | --- | --- | --- | --- |
| **KKT2 S493A** | 41 | 2 (4.9%) | 0 | 1 (2.4%) | **3 (7.3%)** |
| **KKT2 S923A** | 21 | 0 | 0 | 0 | **0** |
| **KKT4 S422A** | 15 | 0 | 0 | 0 | **0** |
| **KKT7 S304A** | 20 | 0 | 0 | 0 | **0** |
| **KKT7 S304S** | 10 | 0 | 0 | 0 | **0** |
| **Total** | 107 | 2 (1.9%) | 0 | 1 (0.9%) | **3 (2.8%)** |

<sup>‡</sup>*Complex mutants refer to mutants that show integration of the repair template in an unexpected way e.g. complete integration on one allele but partial on another, or with mutations not present in the designed repair template.*

### Optimized double-stranded DNA templates improve precision editing efficiency 8-fold

The low efficiency of the ssODN approach in our targets led us to hypothesize that double-stranded DNA (dsDNA) repair templates would be more effective, aligning with other successful selection-free methodologies in kinetoplastids (Vasquez et al., 2018; Kovářová et al., 2022; Asencio et al., 2024; Carnielli et al., 2025b). We systematically optimized the protocol by making two key changes, switching from ssODNs to dsDNA repair templates and increasing the flanking homology arms from 30 nt to 50 bp. Our revised strategy is shown in Figure 1A. We used this strategy to target six phosphosites across KKT2, KKT4, and KKT7 for either phosphodeficient (Alanine), phosphomimetic (Glutamic acid), or synonymous mutations. These sites were the four previously tried with ssODNs, as well as KKT2 S25 and S530. We streamlined our screening process (Figure 1B) by replacing the restriction digest method with a more robust, two-step PCR strategy (Figure 1C). PCRs were completed on parental controls for each primer pair as well as each clone being evaluated.

**Figure 1.**
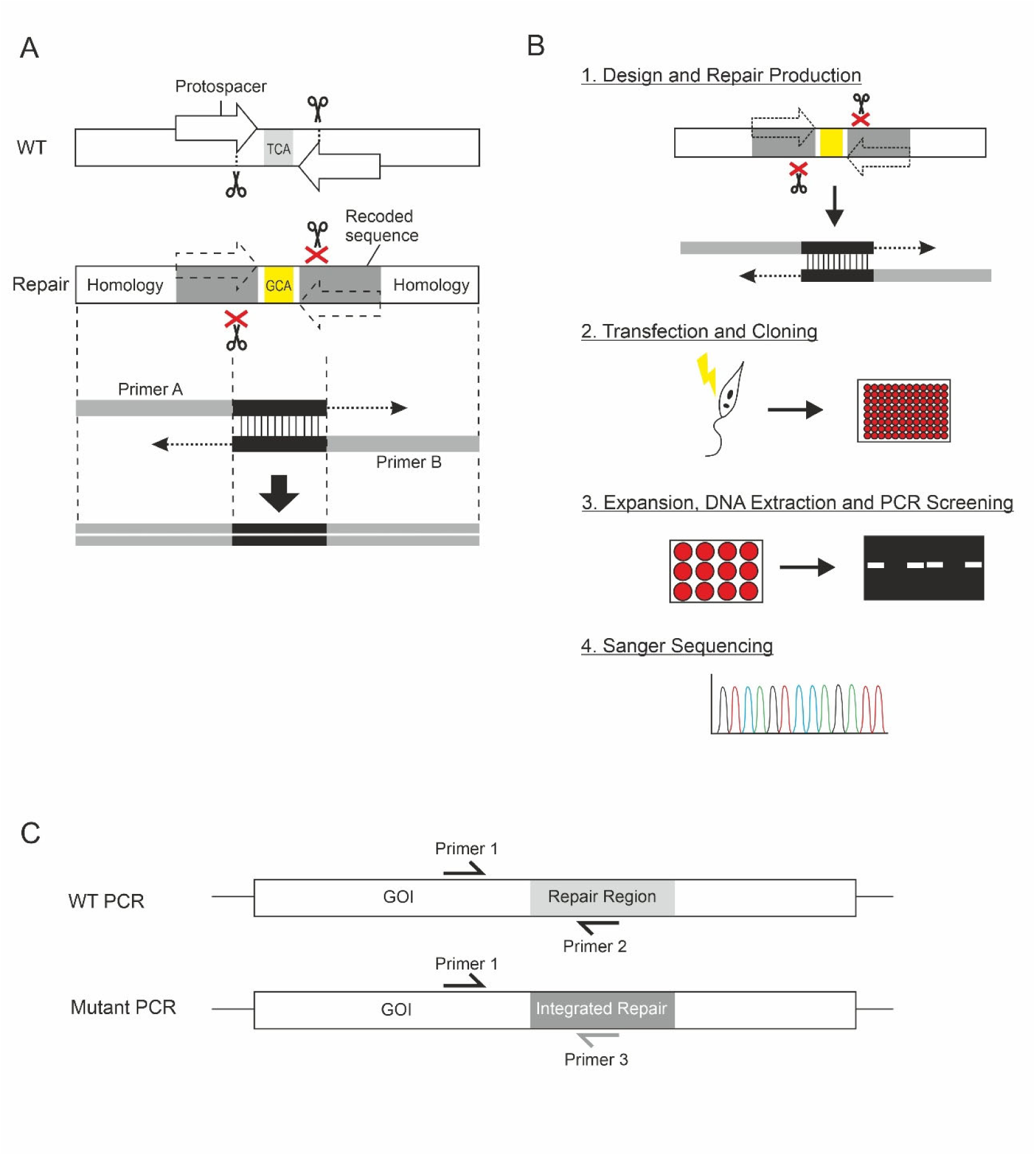
Schematics of the precision editing design and workflow using double-stranded DNA repair templates. A) Repair template design and production. A repair template region is selected and two Cas9 break sites are identified with EuPaGDT (Peng and Tarleton, 2015) – one either side of the target site. The sequence corresponding to the protospacer and PAM sequences are recoded to remove the break sites and provide alternate sequence for screening purposes. The ends of the recoded region are flanked with 50 bp of homologous sequence. After completing the design, primers A and B are synthesised with 19-21 bp of overlapping sequence, which when annealed, is extended by PCR to generate the repair template. B) Precision editing workflow. Part 1 – design and repair template production as in panel A. Part 2 – cells are transfected with the repair template DNA and DNA for expression of the two sgRNAs (Peng and Tarleton, 2015). sgRNA DNA constructs are produced using a PCR method as per Beneke et al. (2017). Following transfection, cells are cloned in 96 well plates and allowed to recover for 1 week. Part 3 – Randomly selected clones are grown in small scale cultures, DNA is extracted and genotype is determined by diagnostic PCRs shown in panel C. Part 4 – clones indicating the presence of a mutant allele by PCR are sent for Sanger sequencing. C) Genotyping screening PCR schematic. Two PCRs are carried out on each clone – one to amplify the WT allele and one to amplify the mutant allele. Both PCRs share a primer in a neutral location outside of the repair template (primer 1), with the second primer that is specific to the genotype (WT – primer 2, mutant – primer 3). Primers 1 and 2 were designed using NCBI Primer BLAST against the WT reference sequence. Primer 3 was designed manually to: have a similar Tm to primer 1, be in roughly the same location in the sequence, and where possible, to have a 3’ base which was different between WT and mutant sequences.

The optimized protocol achieved a dramatic increase in efficiency. We successfully isolated at least one homozygous mutant clone for every target site. Overall, 30.4% of the screened clones showed confirmed integration of the repair template. Most critically, 22.1% of all clones screened were confirmed as homozygous precision mutants by Sanger sequencing (Figure 2 and Supplementary Figure 1). This efficiency represents an order-of-magnitude improvement over the ssODN approach. For instance, the previously failed KKT7 S304A transfection now yielded a successful mutant, while the improved KKT2 S493A transfection produced 9 edited clones out of 12 screened, demonstrating a vast increase in the consistency and success rate of generating single-site mutations. The increased efficiency frequently permits the generation of multiple independent clones carrying the same precision edit, allowing more robust conclusions on target function where a phenotype can be observed.

**Figure 2.**
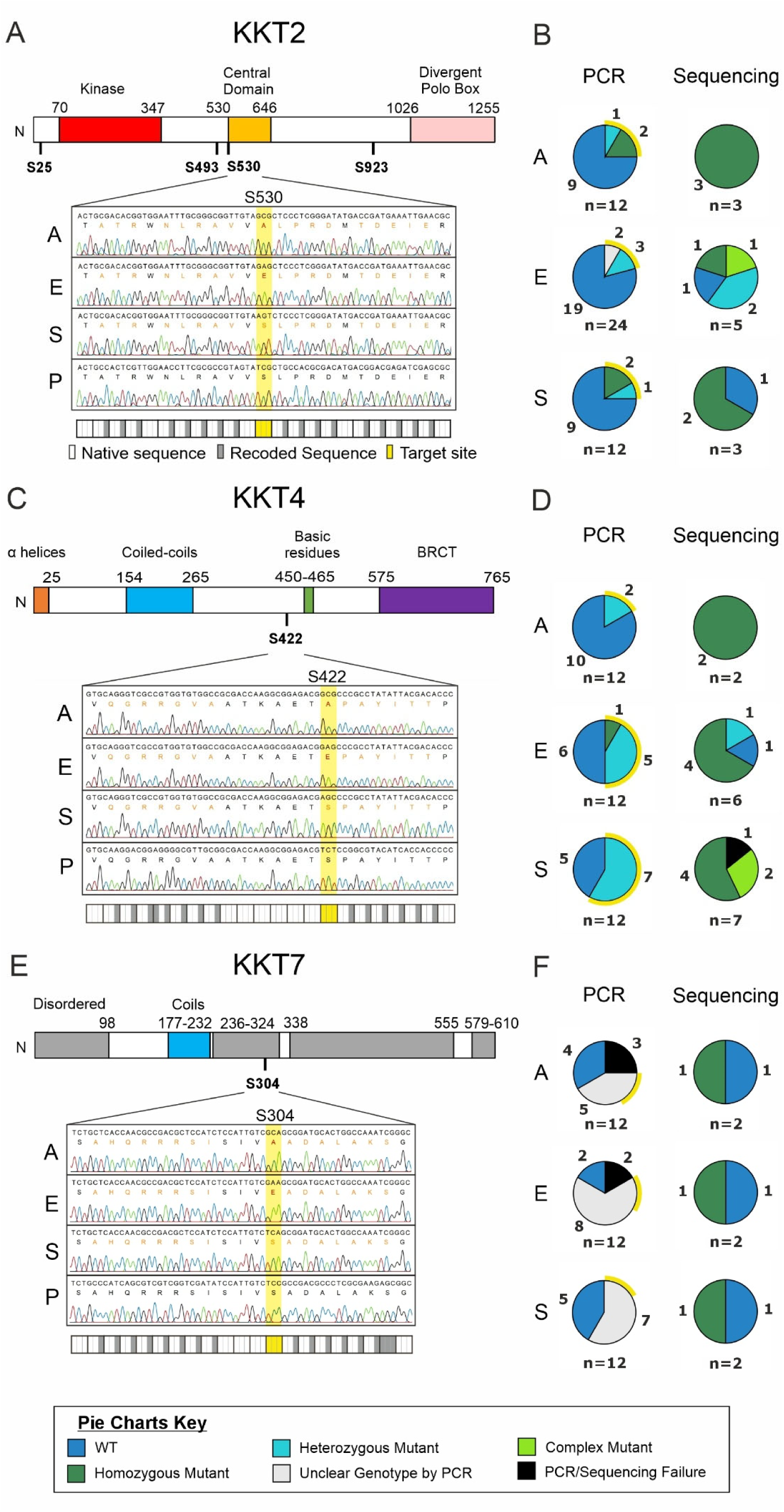
Representative precision editing sequencing and efficiencies. A and B) KKT2 S530 mutants, C and D) KKT4 S422 mutants, and E and F) KKT7 S304 mutants. Panels A, C and E show schematics of KKT protein domains, (Llauró et al., 2018; Marcianò et al., 2021) with target serine phosphorylation sites indicated underneath in bold. The chromatogram from the Sanger sequencing results of one homozygous mutant clone with each mutation as well as the parental sequence is shown below (A – alanine mutant, E – glutamic acid mutant, S – synonymous serine mutant, P – parental T7Cas9 cell line). Mutated bases are indicated with grey rectangles (synonymous mutations) or yellow rectangles (target site mutations) below each set of sequencing. The translation is shown below each DNA sequence, with black text indicating the same protein and DNA sequence as the parental sequence, orange text indicating the same protein sequence but a different DNA sequence to the parental, and red indicating a difference in the protein sequence and hence DNA sequence. Panels B, D and F show the number of clones screened for their respective genotype by PCR, and sequencing of PCR-positive clones. Yellow halos around PCR screening pie charts indicate clones selected for sequencing. A, E and S indicate mutant residue as with panels A, C and E. The number of clones per slice is indicated around the outside of each pie, with the total clones assessed below. For PCR data: WT – PCR product was detected with WT primers but not with primers designed to detect the specific mutation; Heterozygous mutant – PCR product was detected with WT primers and with primers designed to detect the specific mutation, at approximately equivalent intensity; Homozygous – PCR product was detected only with primers designed to detect the specific mutation; Unclear – PCR product was detected with both WT and mutant primers, with either differing intensity in each or additional unknown products; Fail – no PCR product was detected in either reaction. For Sanger sequencing data: WT – both alleles match the parental sequence; Heterozygous – one allele matched the parental sequence, one allele matched the repair template sequence; Homozygous – both alleles match the repair template sequence; Complex – evidence of integration of the repair template but with unexpected mutations; Fail – the sequence was unable to align with either the reference sequence or the repair template sequence.

While most successful clones showed the expected homozygous or heterozygous genotype, sequencing revealed that 5 clones (2.1%) had “complex” integration events. These events highlight the remarkable genomic plasticity of *Leishmania*, where the HDR machinery can incorporate the repair template in unexpected ways. Primarily, we observed complex integration events as asymmetrical integration on each allele, where complete incorporation of the 3’ recoded region occurred on both alleles, but only one allele integrated the 5’ recoded region (seen in KKT4 S422S clones 7 and 12). This was also seen in one KKT2 S493A clone recovered from ssODN transfections. However, we also classed integration events that included secondary mutations as “complex” because of the loss of an adjacent codon (e.g., E496 in KKT2 S493E clone 9) or a point mutation (e.g., C to G in KKT2 S923E clone 22) within the recoded region. Despite these rare complex outcomes, the high overall efficiency of generating the desired homozygous mutants validates the dsDNA template/increased homology approach as the new standard for selection-independent precision editing in *L. mexicana*.

### Single kinetochore phosphosite mutants show minimal phenotypic impact

With an array of genetically defined mutants, we next sought to characterize the role of the individual phosphosites on the cell biology of the parasite. Since our target phosphosites are known to vary in phosphorylation status across the cell cycle (Geoghegan et al., 2022), we hypothesised that mutating them would lead to a phenotypic effect during the cell cycle which could impact growth. We assessed these through an Alamar Blue growth assay and a flow cytometry cell cycle assay. Only one clone, KKT7 S304A clone 9, showed a significantly altered growth rate, displaying an enhanced growth rate (1.8-fold growth of parental at day 5) (Figure 3A). All other phosphodeficient, phosphomimetic, and synonymous mutants grew indistinguishably from the parental T7Cas9 line. Analysis of cell cycle distribution showed that most individual point mutants had no significant difference in the proportion of cells in G1, S, or G2/M phases compared to the parental control (Figure 3B-D).

**Figure 3.**
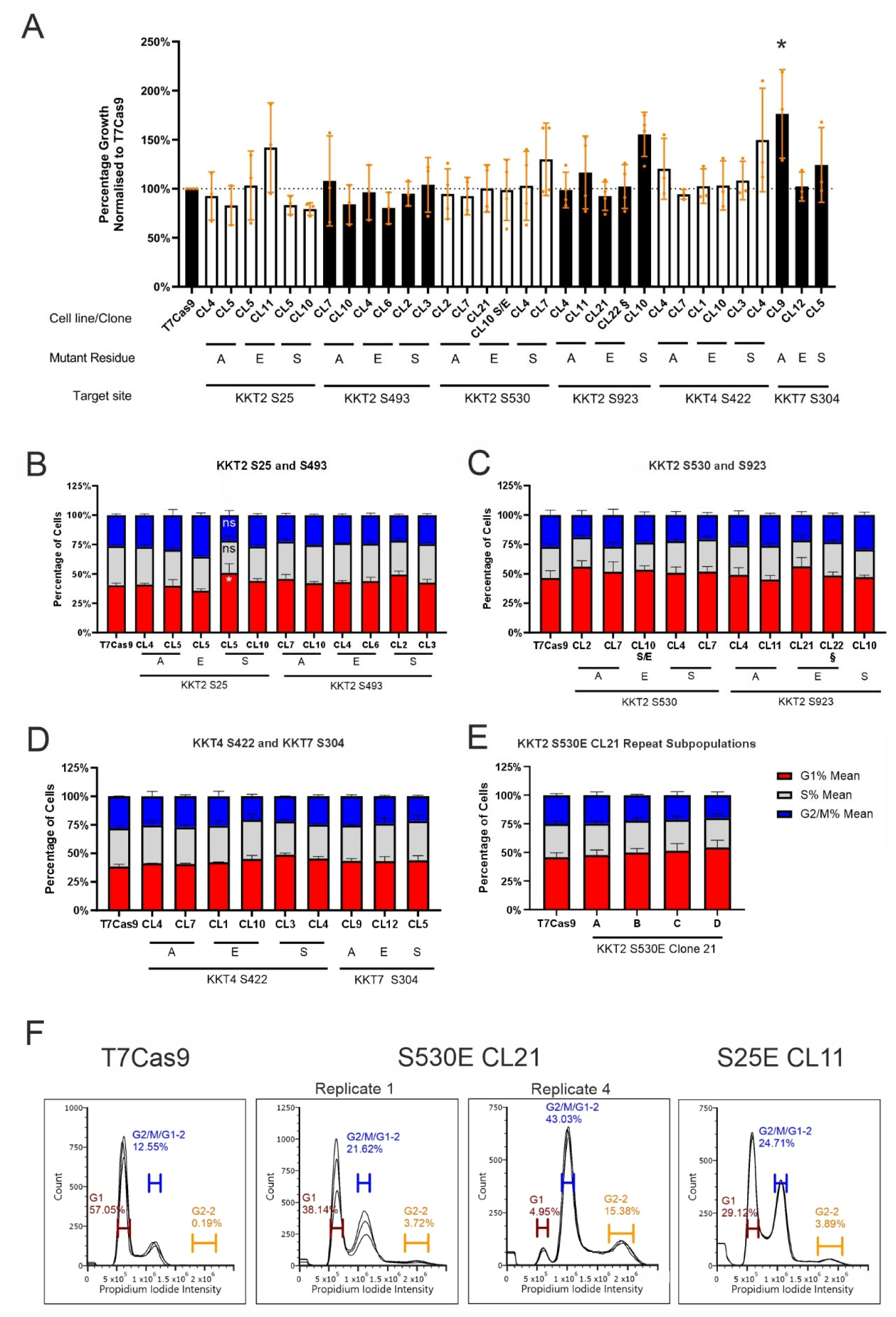
Kinetochore mutant phenotyping. A) Alamar blue growth assay of kinetochore phosphosite mutant clones following 5 days of growth. Colours of bars indicate target site groupings. KKT2 S530E clone 10 is a heterozygote for the S530E mutation, indicated by S/E. KKT2 S923E clone 22 had a heterozygous H927D mutation and is indicated by §. Error bars indicate the standard deviation. * is *p* <0.05. Statistical analysis was completed using a one-way ANOVA test with Dunnett’s multiple comparisons, comparing the means of each cell line with the parental T7Cas9. B-E) Cell cycle analysis of mid-log phase cultures of kinetochore phosphosite mutants assessed by flow cytometry after propidium iodide staining. B) KKT2 S25 and KKT2 S493 mutants, n = 3. C) KKT2 S530 and KKT2 S923 mutants, n = 4. D) KKT4 S422 and KKT7 S304 mutants, n = 2. E) Cell cycle analysis of KKT2 S530E Clone 21 repeat when split into four subpopulations (A to D). n = 3. F) Flow cytometry histograms of propidium iodide intensity for one representative replicate of T7Cas9, the first and last replicate of S530E CL21 and one representative replicate of KKT2 S25E CL11. Each plot shows three technical replicates. Gates used are identical in width between cell lines but have been repositioned to fit the exact intensity of the peaks for each biological replicate. Percentages correspond to one representative technical replicate. In panels B-E, statistical analysis was completed using a 2-way ANOVA test with Dunnett’s multiple comparisons test, comparing each cell cycle stage for each cell line against the corresponding mean of the T7Cas9 parental cell cycle stage. * is *p* <0.05. Even though the percentages are linked (i.e. if G1 is higher, S + G2/M must be lower), each cell cycle stage was assessed independently to simplify the analysis. CL = clone in all panels. Statistical analysis was performed with GraphPad Prism version 8.3.0. Error bars indicate the standard deviation (A) or standard error of mean (B-E).

A subset of phosphomimetic mutants (KKT2 S25E clone 11 and KKT2 S530E clone 21) initially had an uncharacterizable peak during flow cytometry that was consistent with aneuploidy (Figure 3F). However, when KKT2 S530E clone 21 was re-thawed and cultured in independent subpopulations, the aneuploid phenotype was not replicated, and the cells showed a normal cell cycle distribution (Figure 3E). This suggests the aneuploidy was a transient, culture-dependent phenomenon rather than a direct consequence of the genetic edit. Collectively, these phenotypic results indicate that the individual phosphosite mutations generated here do not compromise the essential functions of KKT2, KKT4, or KKT7 under standard *in vitro* growth conditions, suggesting a complex or redundant regulatory mechanism is at play.

### A kinetoplastid Python script standardises design and screening

Leveraging the success of our optimized methodology, we developed a Python script design tool to standardize and accelerate the primer and repair template design process for the community (Figure 4). The script automates the generation of dsDNA repair templates, producing both the desired site-specific mutation (e.g., Serine to Alanine) and a synonymous control mutation for any given target site. It also designs suitable primers for both the repair template production and the final PCR-based clonal screening.

**Figure 4.**
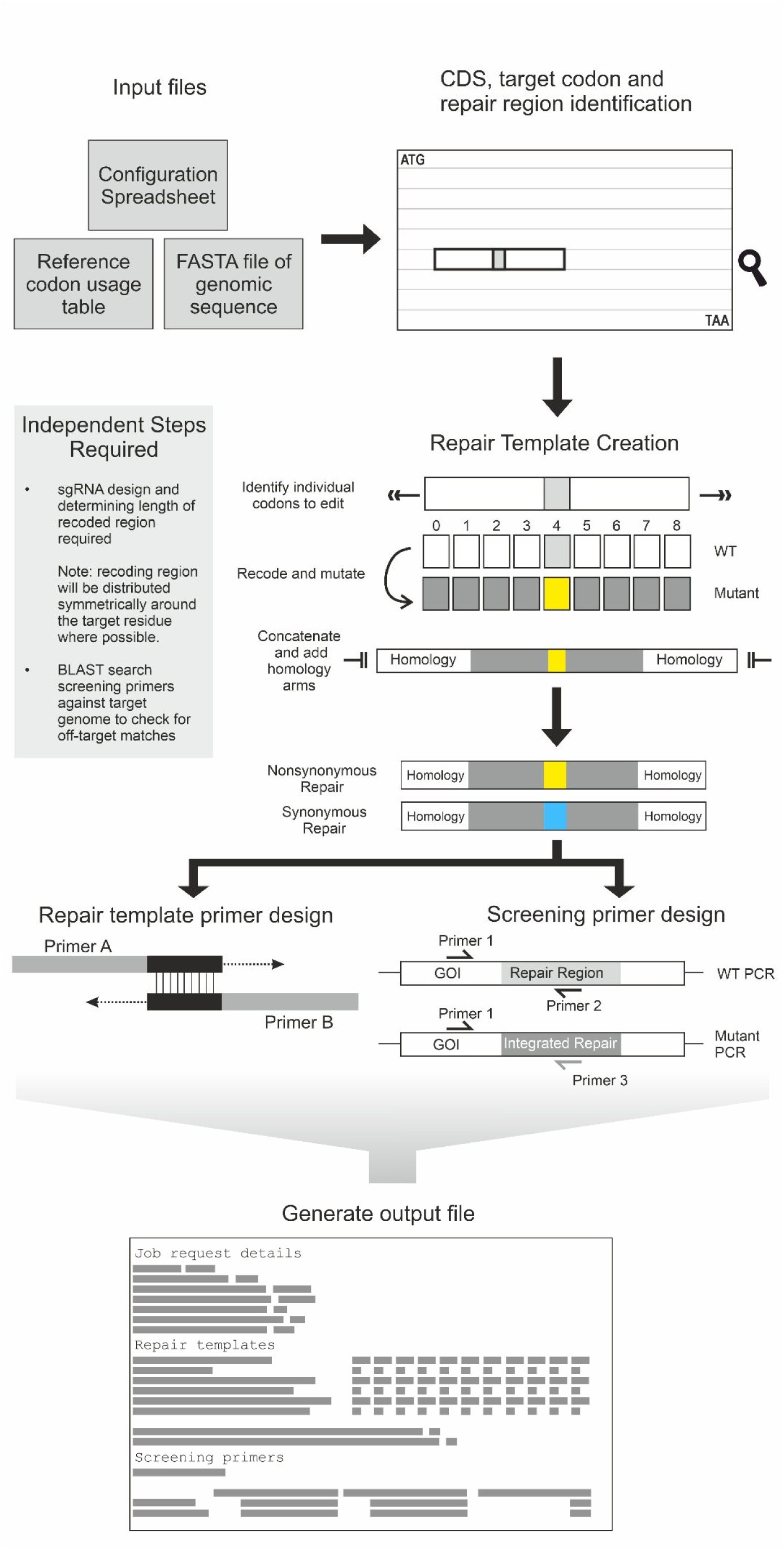
Schematic representation of the Python script workflow. The script requires a FASTA file of the gene of interest, a codon usage table for the organism of interest, and the configuration spreadsheet file specifying the mutation to be generated and the recoding approaches to be used. Initially the script identifies the coding sequence and the target mutation using the information provided in the configuration file. Once a recoding region is identified, each codon within that region is identified, and recoded. The recoding strategy depends on the settings chosen in the configuration file but can be either a harmonised approach (as was used to design the repair templates used in transfections here), using the most or least frequently used codons from the organism, or from a random selection of the alternate codons. A synonymous mutant repair template is also generated for each target, and retains the same synonymous recoding as the target non-synonymous mutant. Once each codon is recoded, the repair template is formed, adding homology arms to each end. Next, primers to generate the repair template are designed using the same strategy as Figure 1A, as well as screening primers using the same strategy as in Figure 1C. Lastly, all the primers and repair templates generated are deposited into an output file.

The tool features an Excel-based configuration file that allows for customized recoding strategies and variable recoding region lengths. The recommended setting for synonymous recoding is called “matched” (utilizes a harmonized recoding to be comparable to native codon usage) and for non-synonymous mutations is “highest” (selects the most frequently used codon). Importantly, the script ensures that the replacement codon for synonymous edits is always distinct from the native sequence, a measure vital for preventing re-cutting by the constitutively active Cas9. Tested successfully on sequences from *L. mexicana* and *T. brucei*, this Python tool provides a rapid, accurate, and species-agnostic solution for the high-throughput design of precision editing constructs in kinetoplastid research.

## Discussion

We have successfully developed an optimized, selection-independent precision editing methodology for *Leishmania mexicana* that uses double-stranded DNA templates with extended homology arms. This protocol dramatically improved editing efficiency over the ssODN approach, yielding 22.1% homozygous precision mutants and demonstrating robust efficacy across multiple essential kinetochore genes. Furthermore, we provide a Python tool to standardize the design of these constructs, offering a “ready-to-go” resource for the kinetoplastid research community.

CRISPR-Cas9 mediated precision editing in kinetoplastids has been a challenging field, primarily due to variable efficiency and a heavy reliance on drug selection markers (Zhang and Matlashewski, 2015; Rico et al., 2018; Wall et al., 2018). While ssODNs have been frequently used, their success has often been tied to drug-based enrichment strategies (Zhang and Matlashewski, 2015; Medeiros et al., 2017; Rico et al., 2018; Wall et al., 2018). In contrast, our group recently demonstrated that ssODNs can be used in a selection-free context to introduce kinase gatekeeper mutations in essential protein kinases, including KKT2. In this context, the approach yielded homozygous mutant clones at a frequency of 11.7% (Carnielli et al., 2025a). However, in the present study, selection-free ssODN-mediated editing resulted in recovery of only 2.8% edited clones across the targeted kinetochore protein phosphosites, suggesting that editing success may be influenced by the nature of the target locus and the specific mutation introduced. To overcome these limitations, we developed an optimized protocol, which utilized 160 bp double-stranded DNA (dsDNA) repair templates with 50 bp homology arms, substantially increasing mutation efficiency in a selection-free setting. These findings are consistent with our other recent work in *L. mexicana*, in which 200 bp dsDNA templates with 48–72 bp homology arms achieved similarly high efficiencies for phosphosite mutation in protein kinase substrates (Carnielli et al., 2025b). When compared to the few published selection-independent studies (Medeiros et al., 2017; Kovářová et al., 2022; Asencio et al., 2024; Carnielli et al., 2025b, 2025a; Novotná et al., 2025), our method is highly competitive, particularly in its simplicity. Our 22.1% homozygous rate compares favourably to the highly variable range reported elsewhere (e.g., 3-30% in other studies, which often only succeeded in specific contexts or organisms) (Medeiros et al., 2017; Kovářová et al., 2022; Asencio et al., 2024; Carnielli et al., 2025a; Novotná et al., 2025). Crucially, our method is effective using the constitutively active T7Cas9 cell line and transient guide delivery and is compatible with the high-throughput CRISPR-Cas9 toolkit established by Beneke et al. (2017). Our method eliminates the laborious step of producing and transfecting recombinant Cas9 ribonucleoprotein particles (RNPs), a requirement for other high-efficiency protocols (Medeiros et al., 2017; Asencio et al., 2024), significantly streamlining the experimental workflow. By demonstrating consistent, high-yield editing without a selection marker, we have established a new, accessible standard for introducing subtle, targeted changes, such as single-site mutations, into the *Leishmania* genome.

One observation from our work is the frequent detection of “complex” mutant genotypes (2.1% of edited clones). These clones exhibited unexpected recombination events, such as partial repair template integration on one allele, the incorporation of unscripted single-nucleotide polymorphisms (SNPs) or small deletions alongside the desired edit. This phenomenon underscores the profound genomic plasticity of *L. mexicana* (Black et al., 2023). Given the absence of the non-homologous end joining (NHEJ) pathway, repair of double-stranded breaks is largely reliant on homology directed repair (HDR) (Passos-Silva et al., 2010). However, the flexibility with which *Leishmania* can incorporate exogenous DNA suggests the involvement of micro-homology-mediated end joining (MMEJ) repair pathway. MMEJ is used more frequently in *Leishmania donovani* than mammalian cells for double-stranded DNA break (DSB) repair, in part because *Leishmania* lack the non-homologous end joining (NHEJ) pathway which is preferred in mammalian cells (Zhang and Matlashewski, 2015, 2019). MMEJ uses short regions of homologous DNA to stimulate repair (5-25 nt) (Zhang and Matlashewski, 2019; Kumari et al., 2025) which aligns with our observation that as little as 11 nt of homology was enough to stimulate partial repair template integration. In order to repair a DSB via the MMEJ pathway, microhomology regions in proximity to the break site are detected by PARP1 and the MRN complex (MRE11, RAD50, and NBS1). PARP1 and the MRN complex resect the DNA until the microhomology sequence is flanking the break, creating single-stranded overhangs in the process (Kumari et al., 2025). Next, these overhanging sequences are removed by endonucleases. Lastly, polymerase theta and the ligase III/XRCC1 complex are recruited to fill in missing sequence and ligate the strands (Kumari et al., 2025). Polymerase theta has already been shown to be essential for *L. donovani* and is required for both MMEJ and single strand annealing DSB repair in the parasite (Zhang and Matlashewski, 2019). MMEJ is often erroneous, so is generally considered to be a back-up repair mechanism, as the resection of the sequence from the break site leads to deletions. However, if the MMEJ pathway was involved in the repair of the DSB breaks in our complex mutants, the presence of the repair template seems to have mitigated loss of sequence in most cases.

We observed complex integration more frequently in repair templates that contained sections of native sequence within the recoded region, such as between the two recoded guide sequences (e.g., KKT4 S422S and the ssODN for KKT2 S493A). This suggests that regions of micro-homology within the template or surrounding genomic sequence may be exploited by the parasite to mediate an aberrant repair. With this in mind, we created the Python script to design repair templates with continuous recoding across the entire recoded region, to avoid opportunities for the parasite to circumvent our intended design. The presence of these complex genotypes, which have not been widely reported in other kinetoplastid precision editing studies, demonstrates that full sequencing of potential mutant clones is paramount to confirm the intended genotype. A deeper mechanistic understanding of this versatile recombination could eventually be manipulated to further enhance or control editing outcomes.

The lack of an apparent growth or cell cycle phenotype in the individual KKT2, KKT4, and KKT7 phosphosite mutants suggests that the regulation of the *Leishmania* kinetochore is not dependent on a single, rate-limiting phosphorylation event, but rather relies on a more intricate, potentially redundant network. This finding aligns with previous studies in *T. brucei*, where disruption of key phosphorylation pathways, such as those involving CLK1/CLK2 kinases, did not always abolish protein function or localisation (Ishii and Akiyoshi, 2020). For example, the absence of phenotype in our KKT2 mutants suggests either that the CLK1 kinase is still able to phosphorylate the critical residues needed for kinetochore assembly, or that assembly in *L. mexicana* is less reliant on this specific event than in *T. brucei* (Saldivia et al., 2021). Further work to generate and test the KKT2 S505/S506 mutants, which correspond to *T. brucei*’s CLK1-targeted sites, using our optimized dsDNA protocol would be a logical next step. The fitness of our KKT7 S304A phosphodeficient mutant mirrors results in *T. brucei* and suggests that while phosphorylation at this site is dynamically regulated during the cell cycle, it is dispensable for basal kinetochore function and cell viability (Ishii and Akiyoshi, 2020). The overall resistance to single-site disruption underscores the robust nature of the parasite’s mitotic machinery and highlights the need for multiplex editing or analysis under environmental stress to reveal the conditional roles of these phosphorylation sites.

The high efficiency of this selection-independent methodology opens a wide range of previously laborious genetic manipulations. Beyond phosphosite analysis, this approach is readily adaptable for inserting small, selection-free epitope tags to study protein localization and interaction without disrupting native UTRs or expression levels. It could also be used to manipulate catalytic residues in enzymes or structural motifs in key proteins to investigate function. Finally, introducing multiple targeted mutations into virulence genes to generate highly stable live-attenuated vaccine candidates should be possible. The high rate of homozygous integration achieved by this method significantly reduces the risk of reversion to the pathogenic wild-type phenotype, a crucial safety consideration for live-attenuated vaccines.

In summary, the optimized dsDNA/increased homology methodology, paired with our automated design tool, represents a significant advance as it provides the necessary throughput and precision to answer complex questions about protein regulation and function in *Leishmania* and other kinetoplastids.

## Methods

### Cell culture

T7Cas9 *Leishmania mexicana* promastigotes (Beneke et al., 2017) were grown in HOMEM media with 10% heat-inactivated Fetal Bovine Serum (FBS) and 1% penicillin-streptomycin (10% FBS HOMEM). T7Cas9 cells were also kept under continual selection with 50 µg/ml hygromycin and 75 µg/ml nourseothricin at 25°C in non-vented TC coated flasks (Corning).

### Design and production of sgRNA and repair templates

Genomic sequences for genes of interest were retrieved from TriTrypDB.org from the *Leishmania mexicana* MHOM/GT/2001/U1103 genome. The target site was identified, and ∼60 nt for single-stranded DNA (ssDNA) repair templates or ∼80 bp for double-stranded DNA (dsDNA) repair templates either side of the target site were selected to create a region of a total of 120 nt (ssDNA) or 160 bp (dsDNA) (Figure 1A). Genomic sequences for this region were used for sgRNA design on EuPaGDT (http://grna.ctegd.uga.edu/) (Peng and Tarleton, 2015). The highest ranking two guides in as close proximity to the target site as possible were chosen – with one making a break before, and the other after the target site. The first and final 30 nt (ssDNA) or 50 bp (dsDNA) of the region were kept as the native sequence to provide the homology arms. Sequences corresponding to the protospacer motifs and PAM sequences were recoded using an alternate codon with the highest frequency of usage from *Leishmania infantum* (from https://www.kazusa.or.jp/codon/). A restriction site addition or removal was also engineered into ssDNA repair templates. The target mutation codon was taken as the highest frequency usage codon for the desired amino acid. A synonymous mutant repair template was also designed to share the same protospacer and PAM site recoding, but selecting an alternate synonymous codon for the target site. sgRNA DNA templates were produced as described previously by Beneke et al. (2017). Single-stranded oligodeoxynucleotide repair templates (ssODNs) were synthesised as oligonucleotides dry (Merck) and resuspended at 2 µg/µl in sterile water. For production of double stranded repair templates two oligonucleotides (Merck) with an overlapping region of 18-20 bp were designed (Figure 1A). Oligonucleotides were annealed and amplified using Q5 polymerase (NEB). Resulting products were confirmed by agarose gel electrophoresis. All oligonucleotide sequences used to generate and validate these cell-lines can be found in Supplementary Table 2.

### Transfection and Cloning

T7Cas9 promastigote cells were grown until mid-log phase. 5 × 10^6^ cells were pelleted at 1000 x *g* for 10 minutes, and the supernatant was removed. Cells were resuspended in 100 µl P3 Primary Cell Nucleofector® Solution (Lonza), appropriate repair template and guide DNA. For single-stranded DNA repair transfections, sgRNA guides were cleaned up using a PCR Purification Kit (Qiagen) prior to transfection using approximately 2.5 µg per transfection, and combined with 10 µg of repair template. For double-stranded repair transfections, repair templates and sgRNA guides were cleaned up using a PCR Purification Kit (Qiagen) prior to transfection, using approximately 5 µg of repair template and sgRNA DNA each per transfection. Cells were electroporated using a Lonza 4D Nucleofector® Unit using programme FI-115, and promptly transferred to pre-warmed HOMEM media containing 20% FBS, 1% penicillin-streptomycin (20% FBS HOMEM), but without addition of other antibiotics. Cells were recovered overnight at 25°C. The following morning, cells were distributed into 96-well plates at a density of 0.5 cells/well in 20% FBS HOMEM (cloning). Double-stranded repair transfections were also supplemented with 10 µM 6-biopterin (Merck) during overnight recovery and cloning. Clones were selected at random after 1 week of further recovery, and passaged into 10% FBS HOMEM for subsequent growth.

### DNA extraction and screening clones transfected with ssDNA repair templates

Genotyping of selected clones was completed through a restriction digest strategy. Stationary phase cells were pelleted at 1000 x *g* for 10 minutes, and washed once in PBS. Pellets were frozen dry at −20°C. After thawing, genomic DNA was extracted using Rapid Extract PCR Kit (PCR Biosystems), as per manufacturer’s instructions, except skipping the addition of water and final centrifugation step.

2 µl of crudely extracted DNA from transfected clones and a T7Cas9 parental cell line control were used as a template for a screening PCR, using VeriFi polymerase mix (PCR Biosystems). This PCR spanned the entire region where the repair template was expected to integrate, as well as some of the surrounding genomic sequence. PCR products were confirmed on an agarose gel. Successful PCR products were purified using a PCR Purification Kit (Qiagen) as per manufacturer’s instructions. Purified PCR products were quantified using a nanodrop and adjusted to the same concentration using the elution buffer.

Due to the inclusion of a restriction site change in the repair template, the genotype could be determined by digesting the previous PCR. 500 ng of purified PCR product from each clone and the parental T7Cas9 cell line was digested with the corresponding enzyme (NEB). The reaction was incubated for 1 hour at the appropriate temperature, and then frozen at −20°C to halt the reaction. Undigested input DNA and digested DNA were run out on agarose gels to determine genotype. Undigested DNA from the parental cells and clones indicating a mutant genotype were sent for Sanger Sequencing (Eurofins).

### DNA extraction and screening clones transfected with dsDNA repair templates

Stationary phase cultures were pelleted at 1000 x *g* for 10 minutes and washed once in PBS. Dry pellets were frozen at −20°C, and DNA was extracted using the Rapid Extract PCR Kit (PCR Biosystems) as per the manufacturer’s instructions, except diluting and centrifuging to remove the debris was not completed. A PCR-based strategy was used to assess the genotype of each clone (Figure 1C). Two PCRs were completed on each clone – one to assess for the presence of a WT allele and the other to assess integration of the repair template. As such, one shared primer was designed to anneal outside of the repair template region. The other primer was designed to anneal inside the repair template region with specificity either to the WT sequence (WT PCR) at that locus or the recoded repair template sequence (mutant PCR).

5 µl of the crudely extracted genomic DNA was used for each screening PCR with VeriFi polymerase mix (PCR Biosystems). Genotype was determined by running the PCR products from each screening PCR (WT and mutant) on an agarose gel. The presence of a band at the expected size was taken to indicate the presence of the respective allele (WT or mutant). Cell lines failing to produce a product in either PCR reaction were designated “PCR fail”. Clones indicating a positive result in the mutant PCR reaction were taken forward for Sanger sequencing. On these clones, an additional PCR that covered the whole repair template was completed with Q5 polymerase (NEB), checked on an agarose gel, and then purified with a PCR purification kit (Qiagen) as per manufacturer’s instructions. PCR products were sent for Sanger Sequencing (Genewiz) with one of the primers used for amplification.

### Alamar Blue Growth Assay

Cultures were grown to mid-log phase in 10% FBS HOMEM, then seeded onto 96-well plates in triplicate with 500 cells per well in 200 µl of 10% FBS HOMEM. The plate was returned to the incubator for 5 days, then supplemented with 40 µl 0.0125% (w/v) resazurin (Alamar blue) in PBS into each well and left to incubate in the dark at 25°C for 4-6 hours. The fluorescence at emission of 590 nm was then measured with a BMG Labtech CLARIOstar® microplate reader. The readings of the wells containing cells were corrected to media-only wells. The mean of the triplicate wells was normalised to the parental T7Cas9 control to calculate the percentage growth.

### Flow Cytometry

Cultures were grown in 10% FBS HOMEM media, and 1 × 10^7^ mid-log phase cells were pelleted at 1000 x *g* for 10 minutes. Cells were washed once in PBS with 5 mM EDTA (PBS-EDTA), and the pellet was resuspended in PBS-EDTA. Cold methanol was added slowly to a final concentration of 70% (v/v) and were left at 4°C to fix overnight. Samples were then diluted to 36.8% methanol (v/v) by adding PBS-EDTA and cells were pelleted as before. The pellet was washed once in PBS-EDTA and resuspended in PBS-EDTA with 10 µg ml^-1^ propidium iodide and 10 µg ml^-1^ RNaseA. Samples were incubated in the dark at 4°C overnight, gently resuspended and transferred to a 96-well plate, splitting the sample between three wells per cell line. Samples were analysed on a CytoFLEX LX355, gating for parasite cells, followed by single cells. Each well was set to record 20,000 events in singles, measuring the propidium iodide, as well as forward and side scatter. FCS Express 7 was used to analyse the results. The gating used to collect the data was replicated for analysis, and the number of cells was plotted against the propidium iodide intensity. The proportion of cells under each peak was assessed using the built-in DNA content analysis (Multicycle) to fit 1 cycle using model 5. The percentages of cells in each cell cycle stage (G1, S and G2/M) were collated for the triplicate wells, which was then averaged. The replicates were then averaged and plotted.

### Python Script Development

The Python script was created with Python 3.10.9 as well as the additional packages in Table 2 (Koressaar and Remm, 2007; Cock et al., 2009; Mckinney, 2010; Untergasser et al., 2012; Harris et al., 2020).

**Table 2.**
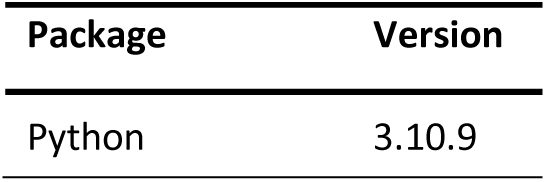
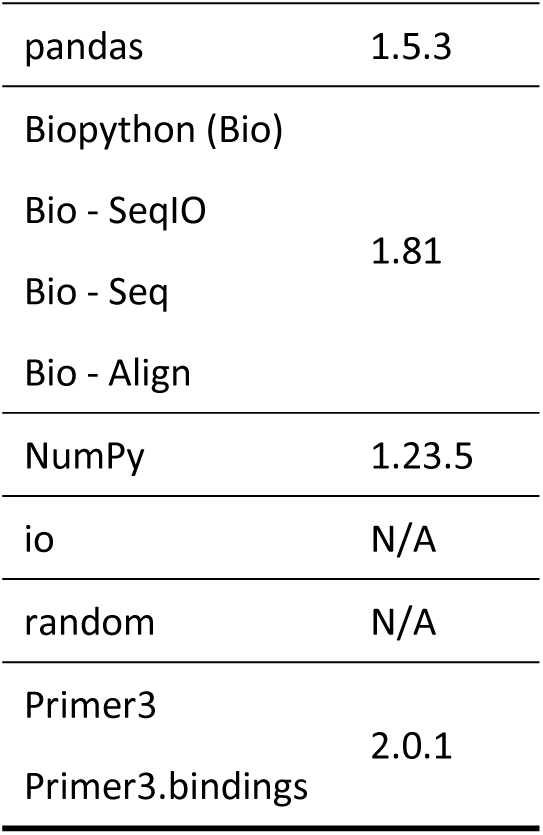
para.

Briefly, the user supplies a FASTA file of their gene sequence, a codon usage table, and the Excel configuration file to the script. The script identifies the target codon to mutate and the surrounding region up to the specified length for recoding, centring this region over the target codon. Then, the script recodes the region using the selected strategy. There are four types of recoding the script can perform. The “matched” setting can only be applied to synonymous mutations, but the other three (“random”, “highest” and “lowest”) can be applied to both synonymous and nonsynonymous mutations. “Random” is as the name suggests, a random unbiased selection of possible codons for the desired amino acid using the random Python package. Unlike the other recoding methods, “random” will cause a different output repair sequence on each execution of the code when given the same inputs (excluding methionine, tryptophan and amino acids with only two possible codons when performing synonymous mutations). “Highest” and “lowest” settings are in reference to the frequency usage of the codon sequences. The user supplies a codon usage table as part of the required inputs (see Figure 4) which will dictate which codon is selected for each amino acid, with “highest” referring to the most used codon, and “lowest” the least used. These allow the user to bias their recoding to use more common or rarer codon sequences, as desired. “Matched” is essentially a harmonized codon selection - choosing the codon that is most similarly used compared to the input codon and is akin to the design strategy used in the *in vitro* experiments. When the chosen recoding method is applied to nonsynonymous mutations, all codon sequences for the respective amino acid are considered in the selection process. However, when the recoding method is applied to synonymous mutations, the WT codon sequence is removed from the available codons to choose from to ensure a mutation occurs (except for methionine and tryptophan codons).

Following repair template design, the script designs screening primers and primers to create the repair template. Both use the Primer3 package to identify suitable annealing sequences (Koressaar and Remm, 2007; Untergasser et al., 2012). Briefly, in the case of repair template primers, Primer3 is used to identify a region to use to anneal the two halves of the repair template together to allow amplification such that each primer is no longer than 120 nt. Due to the small input sequence used for this, primer design settings are quite relaxed (See Supplementary Table 3 for details). For screening primer design, Primer3 initially designs a primer set using the WT sequence, such that one primer is upstream of the repair template and the other within the region that is recoded in the repair template generated. This is the WT screening primer pair. Subsequently, the upstream primer is retained and used to design a complementary primer within the recoding region of the repair template. This is the mutant primer screening pair. As the screening primers are only designed using the input genomic sequence, their specificity may be limited as they are not BLAST searched against the reference genome for the organism.

The script has been tested on a range of sequences from both *Leishmania mexicana* and *Trypanosoma brucei* to confirm the correct mutations have taken place and that primer sequences selected are real sequences. The script has been deposited on Github and is archived on Zenodo (DIO 10.5281/zenodo.18155504).

---

**Supplementary Figure 1.**
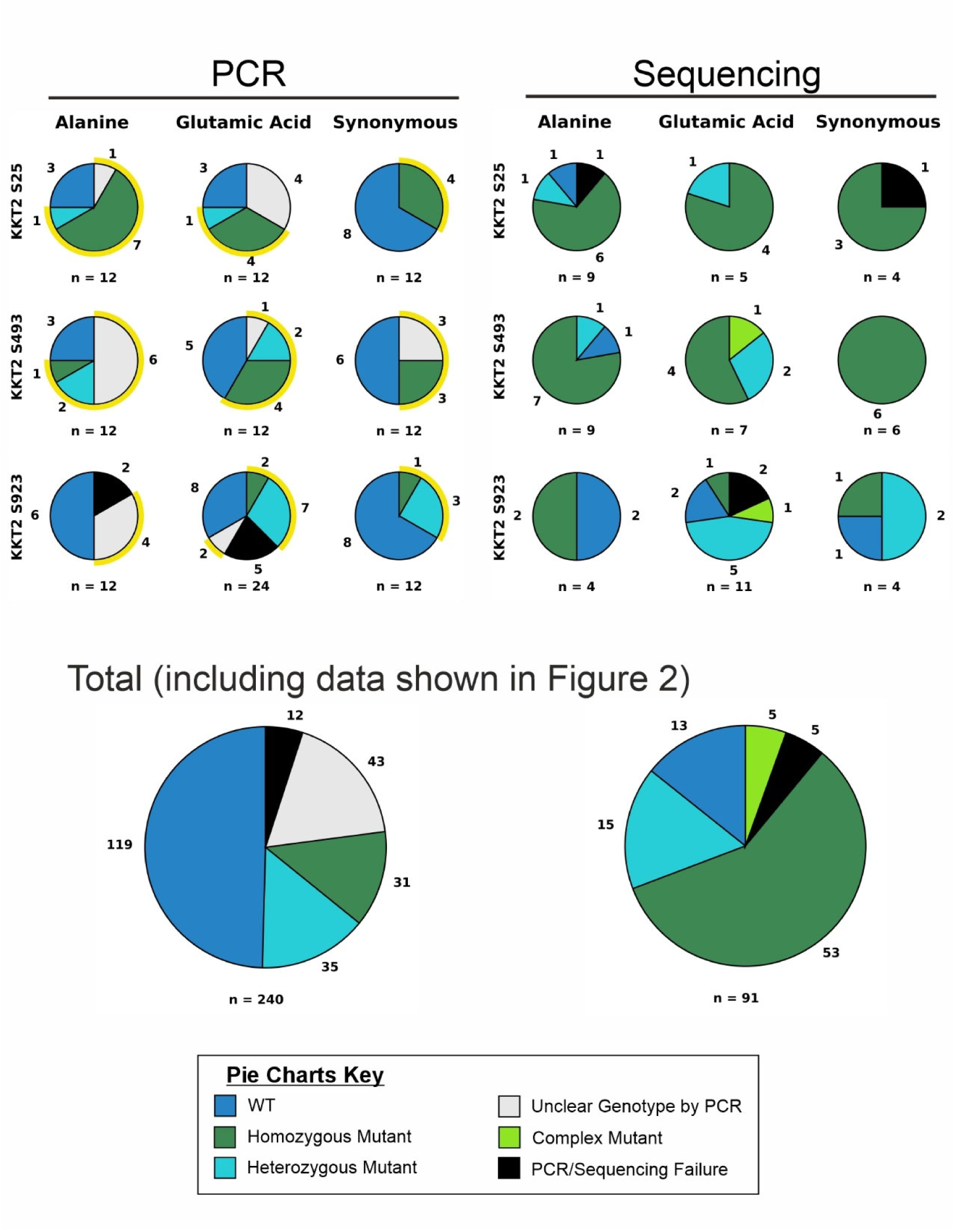
Additional target mutations generated with the precision editing methodology with their genotypes as detected by PCR and Sanger sequencing. The number of clones per slice is indicated around the outside of each pie, with the total clones assessed below. Yellow halos around PCR screening pie charts indicate clones selected for sequencing. The combined data for all mutations generated for both PCR screening and Sanger sequencing is shown at the bottom. For PCR data: WT – PCR product was detected in the WT primer set reaction and not in the mutant set reaction; Heterozygous – PCR product was detected in both WT and mutant primer set reactions with approximately equivalent intensity; Homozygous – PCR product was only detected in mutant primer set reaction; Unclear – PCR product was detected in both WT and mutant primer sets with either differing intensity in each or additional unknown products; Fail – no PCR product was detected in either reaction. For Sanger sequencing data: WT – both alleles match the reference sequence; Heterozygous – one allele matched the reference sequence, one allele matched the repair template sequence (identified by dual peaks of similar height in the chromatogram); Homozygous – both alleles match the repair template sequence; Complex – evidence of integration of the repair template either to different extents on each allele, or with unexpected mutations; Fail – the sequence was unable to align with either the reference sequence or the repair template sequence.

**Supplementary Table 1.**
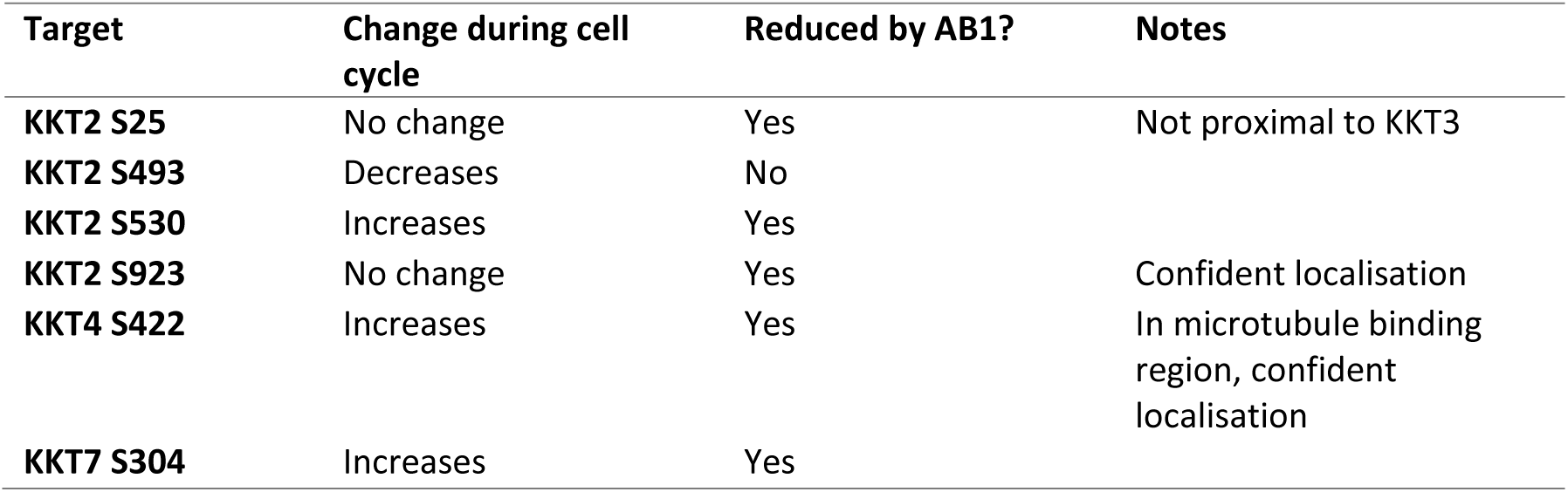
Kinetochore phosphosite data from Geoghegan et al. (2022) used to determine target sites of interest.

**Supplementary Table 2.**
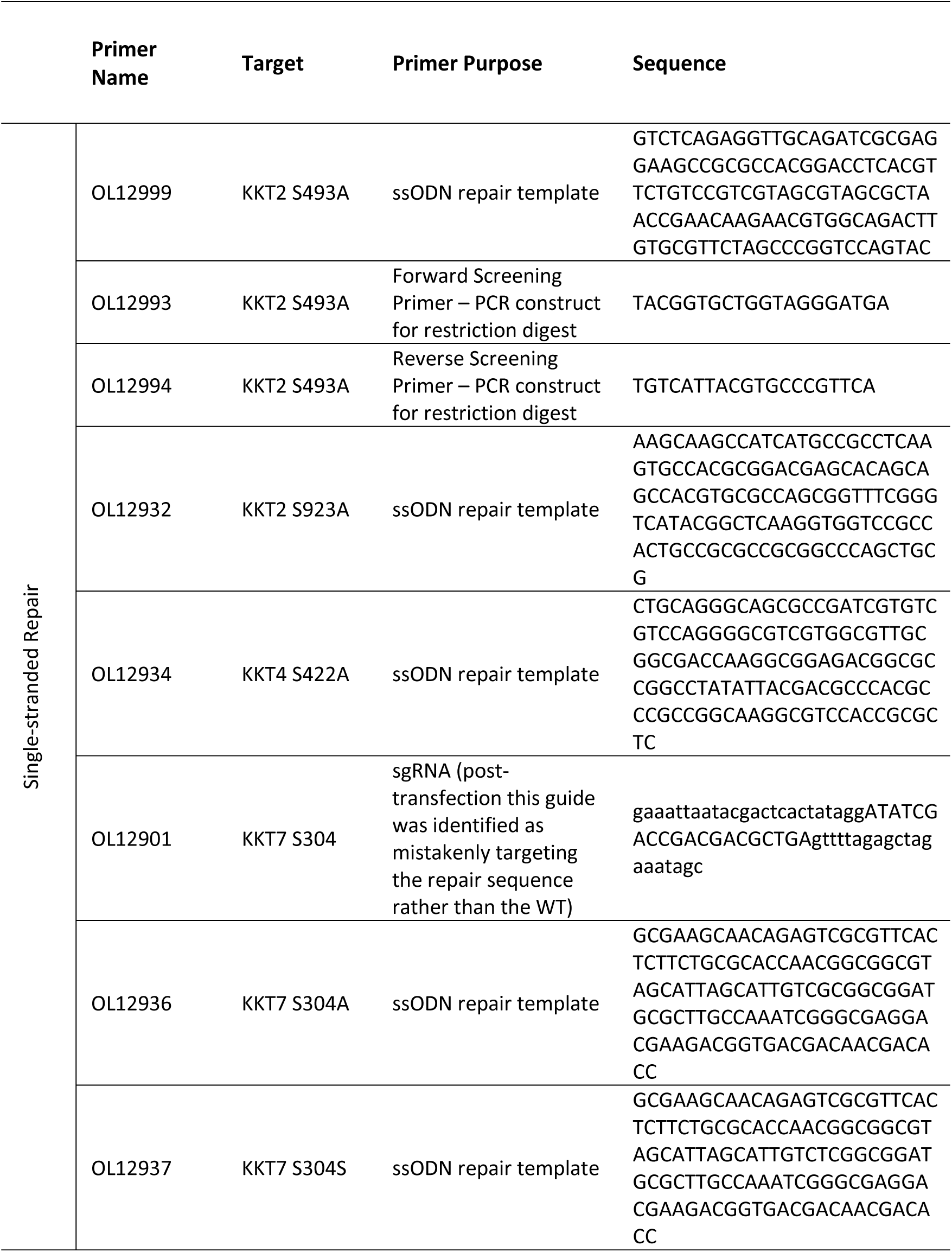
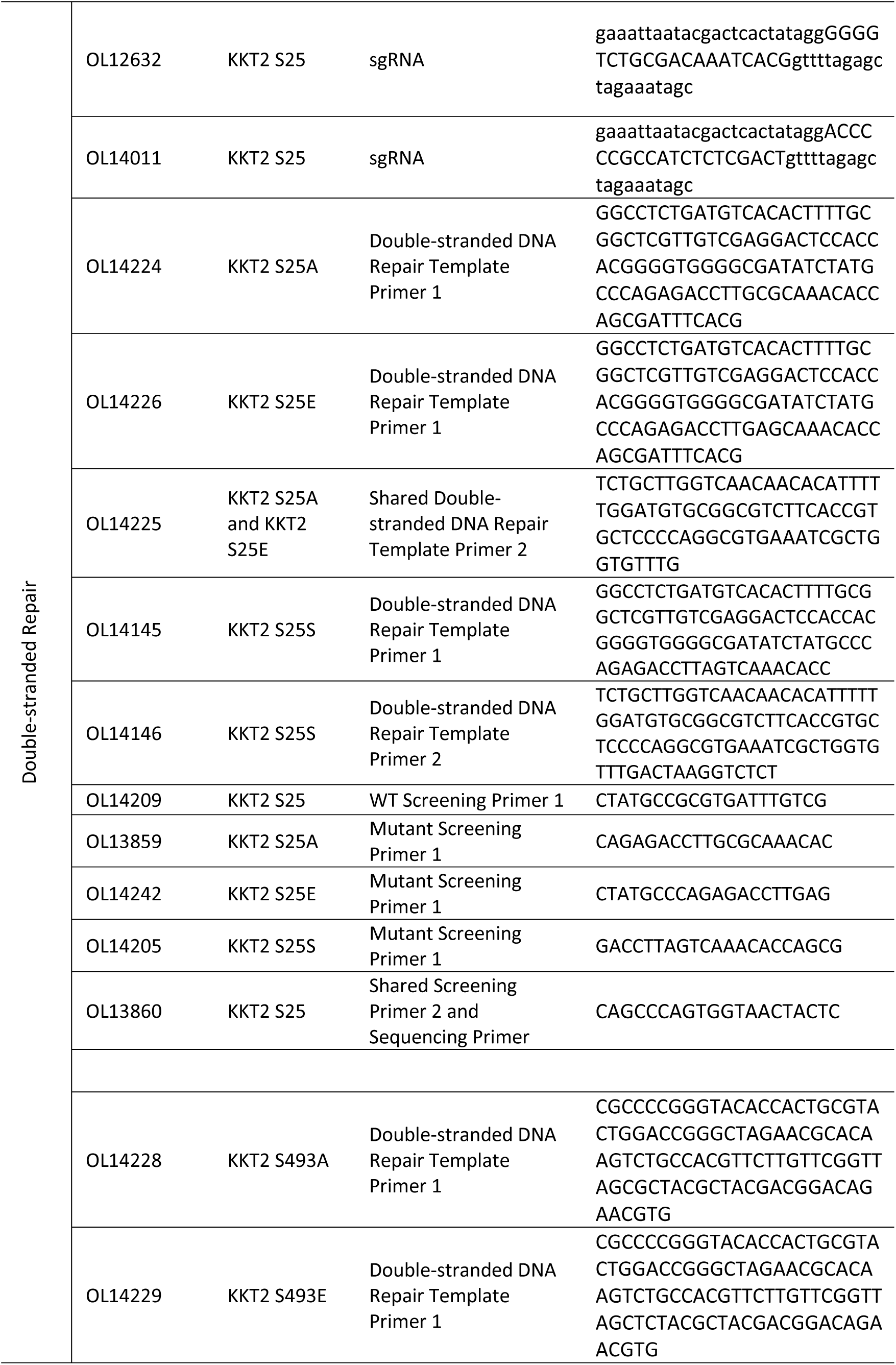
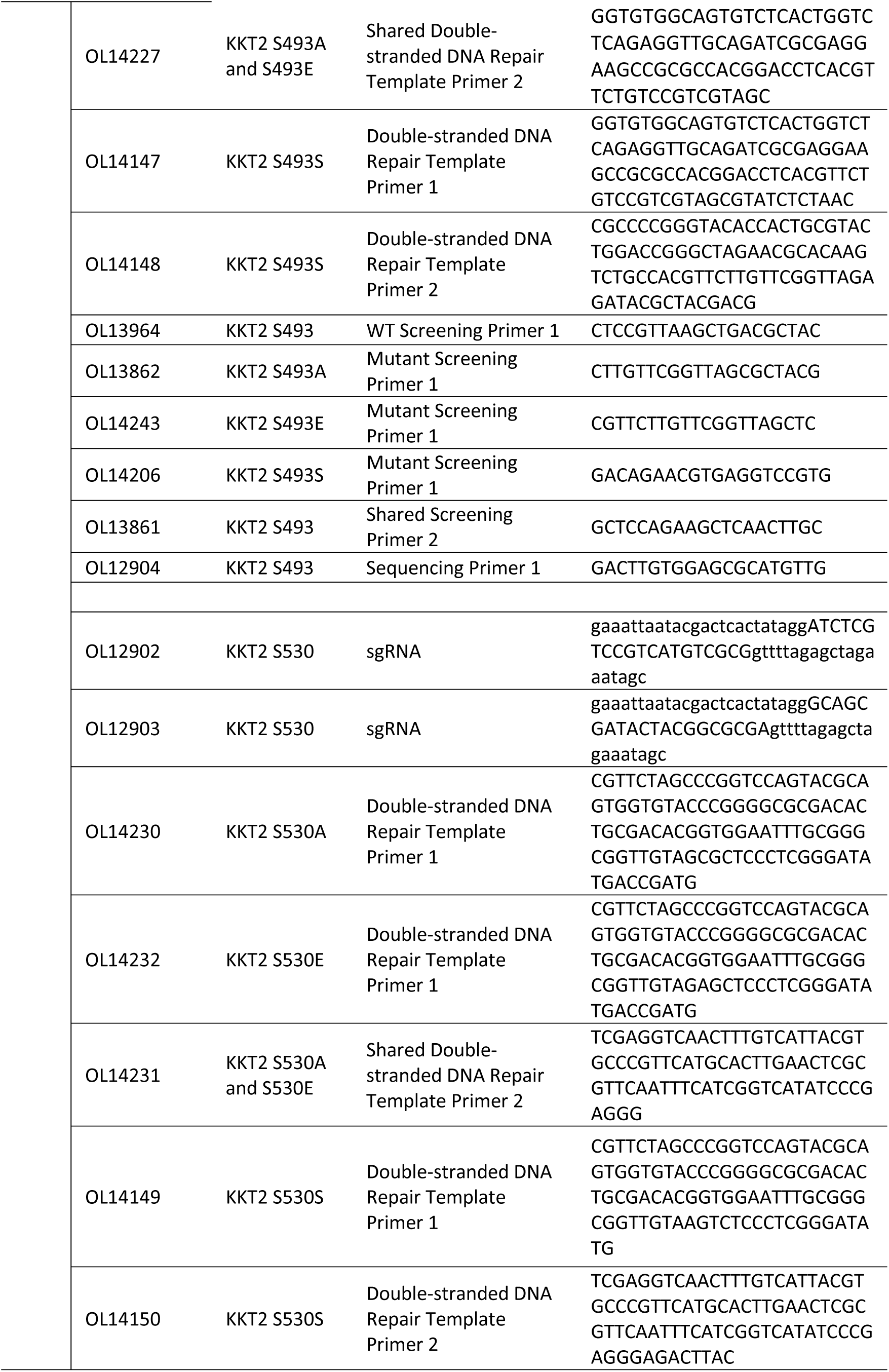
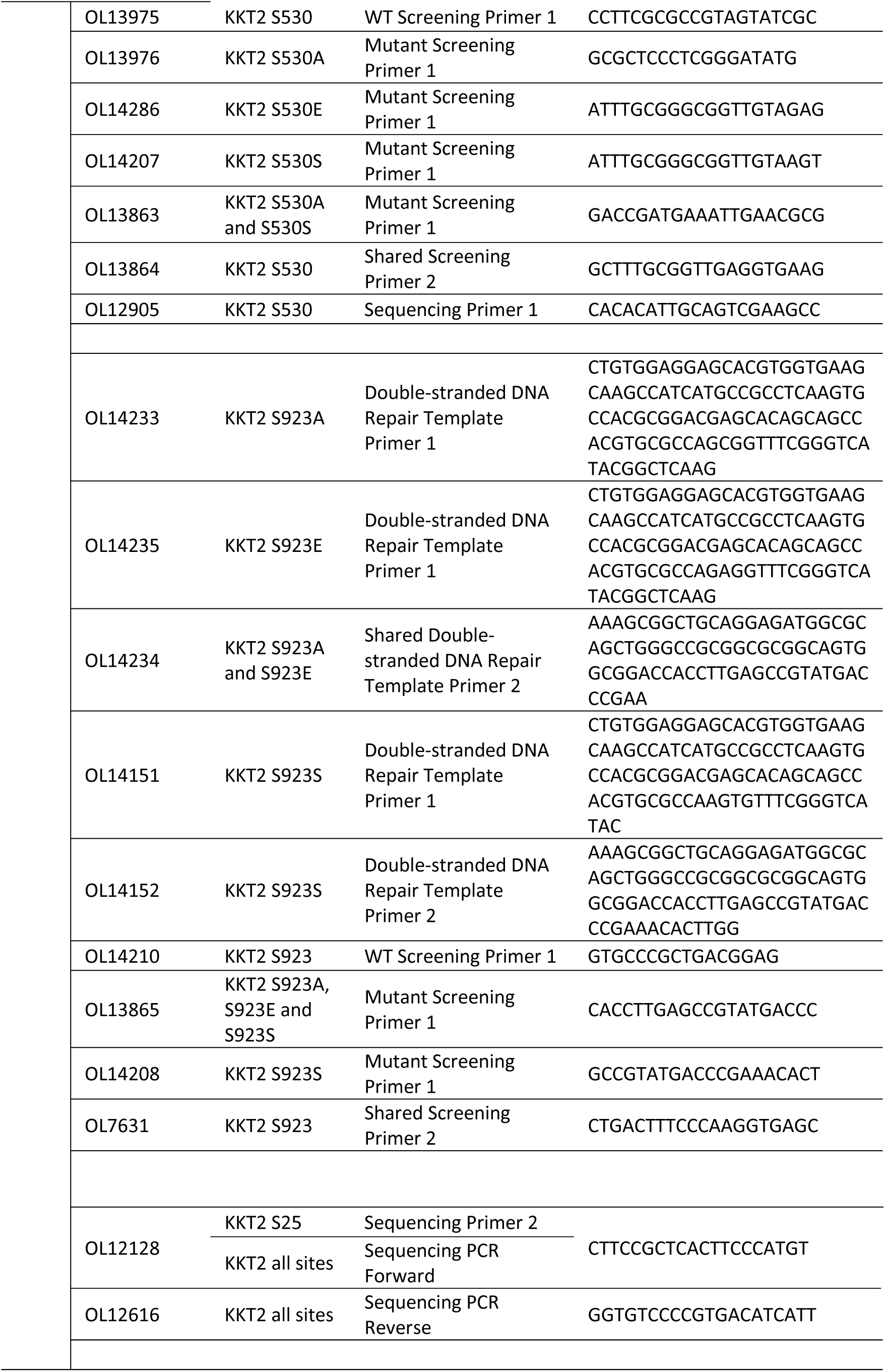
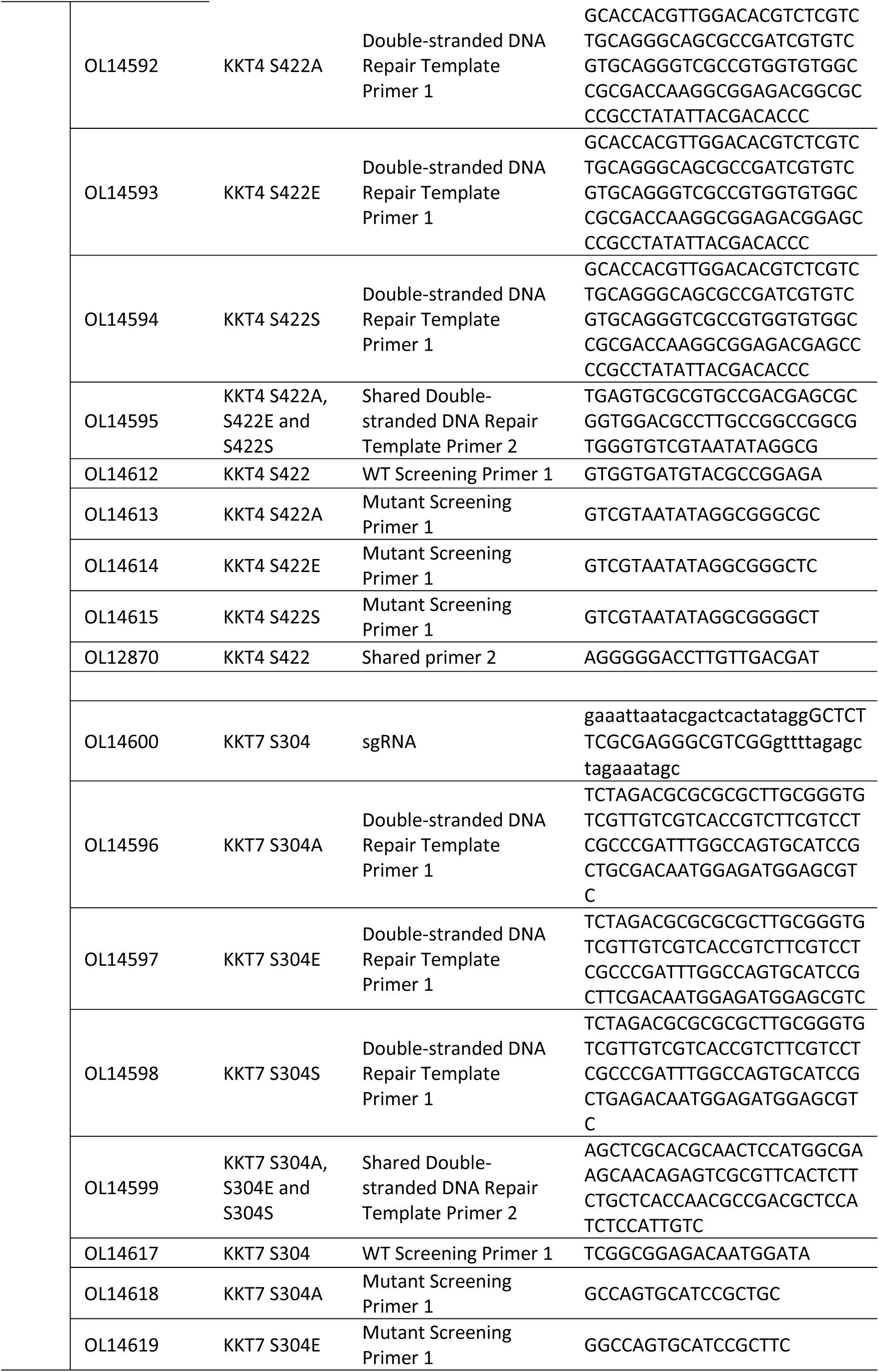
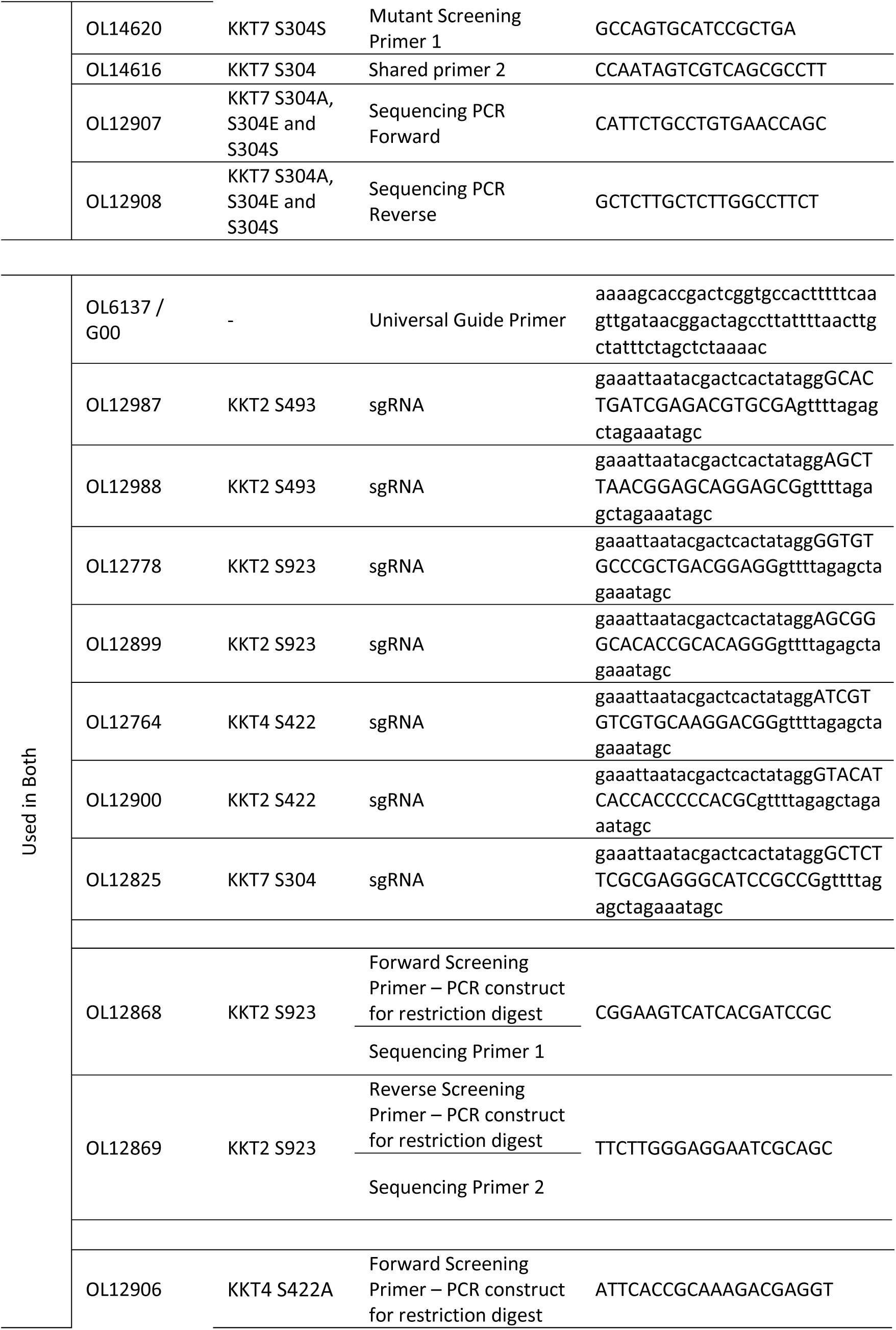
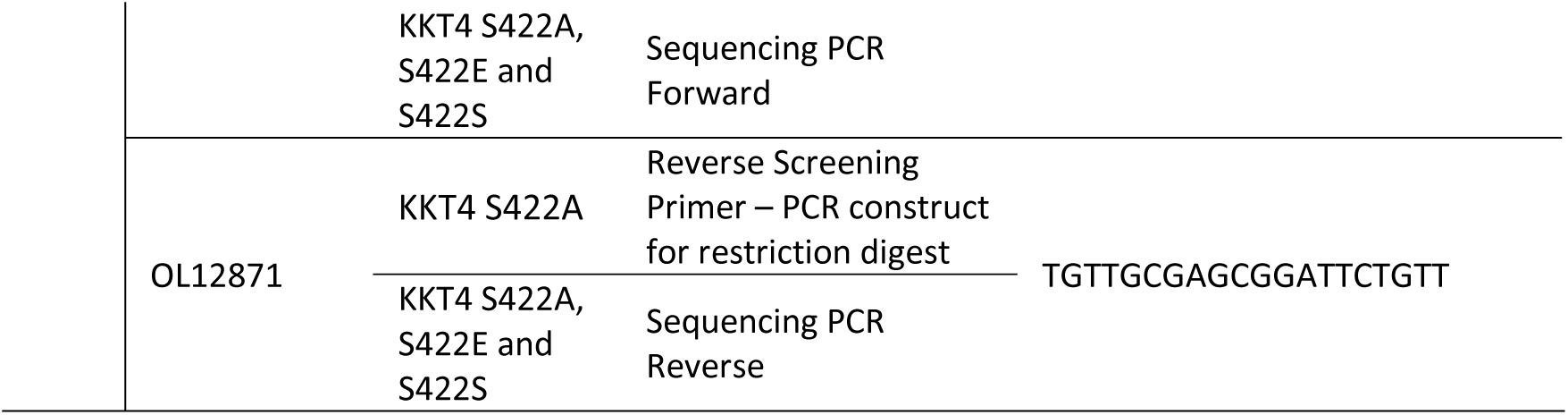
Oligonucleotide sequences used in this study. All sequences are provided in the 5’ to 3’ orientation.

**Supplementary Table 3.**
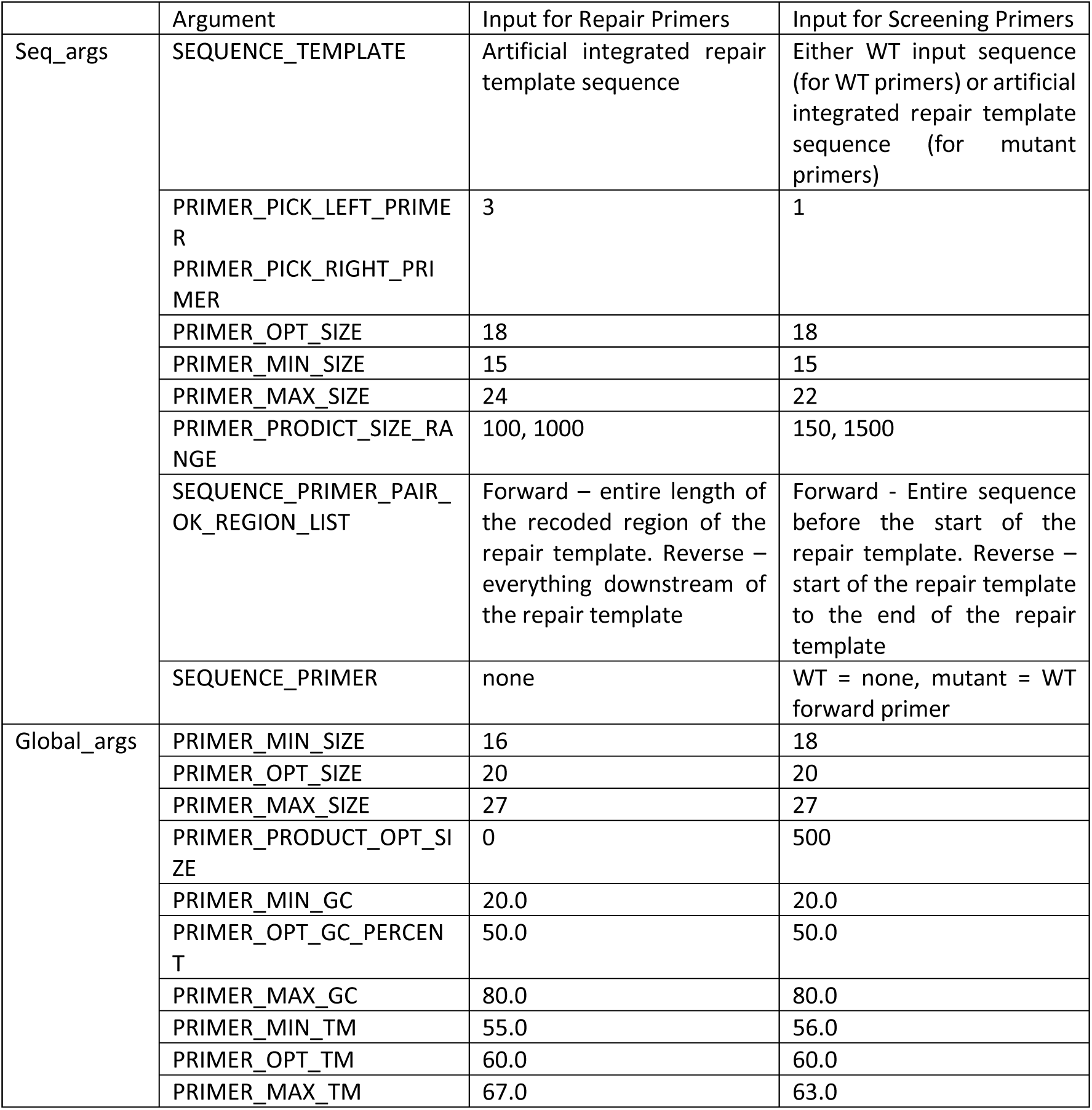
Key Primer3 settings used in the Python script. Other input settings required were the suggested settings based on the Primer3 documentation.

## Notes

### Competing Interest Statement

The authors have declared no competing interest.

### Funding

- Wellcome Trust: 223045/Z/21/Z

